# Fish Blood Response to Ash-Induced Environmental Alkalinization, and their Implications to Wildfire-Scarred Watersheds

**DOI:** 10.1101/2024.01.05.574400

**Authors:** Garfield T. Kwan, Trystan Sanders, Sammuel Huang, Kristen Kilaghbian, Cameron Sam, Junhan Wang, Kelly Weihrauch, Rod W. Wilson, Nann A. Fangue

## Abstract

Changes in land use, warming climate and increased drought have amplified wildfire frequency and magnitude globally. Ash mixing into aquatic systems after wildfires rapidly increases water pH, creating an additional threat to wildlife, especially species that are already threatened, endangered and/or migratory. Here, Chinook salmon (*Oncorhynchus tshawytscha*) yearlings acclimated to 15 or 20°C were exposed to an environmentally relevant concentration of ash (0.25% w/v) which caused water pH to rapidly rise from ∼8.1 to ∼9.2. Mortalities occurred within the first 12 hours, and was higher at the higher temperature (33 versus 20 %). The greatest differences in blood chemistry between the two temperatures were dramatically greater (∼7.5-fold) and very rapid (within 1 hour) spikes in both plasma total ammonia (to ∼1200 µM) and lactate (to ∼6 mM) in warm-acclimated salmon, whereas cold-acclimated salmon experienced a much smaller and gradual rise in plasma total ammonia. Salmon at both temperatures experienced extracellular and intracellular alkalosis within 1 hour that recovered within 24 hours, but the alkalosis was smaller in magnitude in fish at warmer temperature. Impacts on plasma ion concentrations were relatively mild and plasma glucose increased by 2- to 4-fold at both temperatures. Notably, the increase in plasma total ammonia in fish at the warmer temperature was far faster and much greater than those reported in previous studies exposing fish despite higher water pH (9.4-10.5) induced without using ash. This suggests that ash has physiological impacts that cannot be explained by high water pH alone which may relate to the complex mixture of metals and organic compounds also released from ash. This demonstrates post-wildfire ash input can induce lethal yet previously unexplored physiological disturbances in fish and highlights the complex interaction with warmer temperatures typical of wildfire-scarred landscapes.

## Introduction

Wildfires (controlled, cultural, and natural) are known to aid in pest removal and native plant reproduction (Pausas and Keeley, 2019), and are essential for healthy Mediterranean climate ecosystems (Syphard et al., 2007). The frequency and magnitude of wildfires across the world have increased, and this is attributed to anthropogenic factors including changes in land use, warming climate, and increased droughts (Westerling et al., 2006; Huang et al., 2015; Pausas and Keeley, 2021). In the aftermath of wildfires, subsequent precipitation (snowmelt, rainstorms) can move ash across the charred watershed, and ultimately increase the concentration of sediments, trace elements, pollutants, and fire retardants in aquatic systems (Adams and Simmons, 1999; Giménez et al., 2004; Costa et al., 2014; Burton et al., 2016; Emelko et al., 2016; Raoelison et al., 2023). Adverse impacts to diverse aquatic organisms have been reported and include slower development, behavioral change, and shifts in food web dynamics (Spencer et al., 2003; Wells et al., 2004; Beganyi and Batzer, 2011; Nunes et al., 2017; Gonino et al., 2019; Gomez Isaza et al., 2022; Muñiz González et al., 2023).

Wildfires are also known to alter the pH of freshwater systems. The ash-alkaline hypothesis proposes post-wildfire ash input releases alkaline elements (anions) into aquatic systems, which in principle should increase both pH and alkalinity levels (Bayley and Schindler, 1991). This impact on the environment has been observed in many wildfire studies, and its impact can be observed lasting ∼5 years (Paul et al., 2022). For instance, one month after the 2012 High Park fire near Fort Collins (Colorado, United States), the Cache la Poudre River water pH had risen from ∼7.9 to ∼8.5 (Son et al., 2015). In another instance, two years after the 2007 Angora Fire at Lake Tahoe (California, United States), the pH of Angora Creek in the unburned forest was ∼6.25, whereas the pH at the site of the wildfire and downstream remained elevated at ∼7.0 (Oliver et al., 2012). Moreover, algal growth in freshwater systems is stimulated by greater seasonal light availability and increased nutrients as ash is washed into the system (Spencer et al., 2003; Robson et al., 2018), and this can result in dramatic diurnal swings in water pH (Sherson et al., 2015; Kwan and Lehmann, *unpublished data*). Other factors (e.g. baseline water pH, rainfall, flow rate, soil pH, wildfire burn intensity) also influence these dynamic aquatic systems, and altogether these dramatic changes in water pH would undoubtedly challenge the acid-base regulatory capacities of aquatic organisms.

Current United States Environmental Protection Agency’s Water Quality Criteria deems freshwater pH to be between 6.5 to 9.0 to be suitable for aquatic life. Although the above examples do not exceed these values, there are many aquatic systems with naturally high pH averaging at ∼8.0 (e.g. Putah Creek near Davis CA; Roaring Fork River near Glenwood Springs, CO; Snake River near Twin Falls, ID; Mississippi River near Fulton, IL), ∼8.5 (e.g. Green River, UT; Lake Mattamuskeet, NC), to ∼9.0 (e.g. Upper Klamath River, OR; Truckee River and Pyramid Lake, NV) (U.S. Geological Survey, 2016; Pyramid Lake Paiute Tribe, 2019) that are near to or already exceeding the upper limit of the recommended pH range (U.S. Environmental Protection Agency., 2013). Not only are these systems more susceptible to the alkalinizing impacts of post-wildfire ash-input, but they are also home to a variety of threatened and endangered species and could complicate their conservation management. Some of these organisms include the endangered coho salmon (*Oncorhynchus kisutch*) and threatened steelhead trout (*Oncorhynchus mykiss*) at Scott Creek, CA, the threatened Chinook salmon (*Oncorhynchus tshawytscha*) at Putah Creek, CA, the threatened green sturgeon (*Acipenser medirostris*) at Klamath River, OR, and the threatened Lahontan cutthroat trout (*Oncorhychus clarki henshawi*) at Truckee River and Pyramid Lake, NV). Moreover, anadromous fishes with fixed reproductive timelines such as salmon species and steelhead trout (many of which are threatened or endangered) may be among the most vulnerable as the required ionic, osmotic, and acid-base (IOA-B) regulation to mitigate ash-induced alkalinization could further aggravate the exhausted spawning adults returning from the ocean and/or compromise their eggs and larval offspring by challenging their internal pH and NH_3_ regulation (see below). Furthermore, fishes living in low alkalinity conditions (e.g. ∼9 μmol/kg at Lake Notasha in OR, United States) (Stoddard, 1987; Eilers et al., 1990; Catalan and Camarero, 1993; Clow et al., 1996) are more vulnerable as ash-induced alkalization as water pH can be more greatly affected. Finally, while this paper focused primarily on the United States, other areas of around the world such as Canada (Reavie and Smol, 2001) and China (Wang et al., 2003) also have high pH systems that could be influenced by wildfire-ash input.

Despite over a century of investigation and many excellent reviews detailing ionic, osmotic, and acid-base regulation (IOA-B) in teleost fish (Claiborne and Heisler, 1984; Cameron, 1989; Claiborne et al., 2002; Evans et al., 2005; Marshall and Grosell, 2006; Tresguerres et al., 2023), only a handful of studies have investigated blood pH and acid-base response to environmental alkalosis (Wilkie and Wood, 1991; Hemming and Hanson, 1992; Wilkie et al., 1993; McGeer and Eddy, 1998; Scott et al., 2005; Mcgeer et al., 2011). When freshwater teleosts are exposed to high pH conditions, there is an immediate increase in blood pH (pH_e_) (Wilkie and Wood, 1991; Wilkie et al., 1993). According to the classic Davenport acid-base physiology, fish must quickly compensate for blood alkalosis by simultaneously accumulating H^+^ in their blood and expelling HCO_3_^−^ into the water, presumably through their gill ionocytes. Rainbow trout (*Oncorhynchus mykiss*) exposed to pH 9.5 water were able to stabilize their pH_e_ within 8-24 hours of exposure (Wilkie and Wood, 1991), and the few mortalities observed were associated with cannulation procedures rather than the acid-base stress. In contrast, Lahontan cutthroat trout, which resides in pH 9.4 at Pyramid Lake (Nevada, United States), were exposed to pH 10 but were unable to recover their pH_e_ and >50% died after 72 hours of exposure (Wilkie et al., 1993). In addition, the high pH exposure and subsequent rise in blood pH_e_ reduce [H^+^] and hinders the process of ammonia excretion as increased proportion of gaseous NH_3_ (compared to ionic NH_4_^+^) in the environment slows the outward diffusion of NH_3_ as well as net total ammonia excretion rate, which ultimately results in a rapid rise in blood total ammonia (Wilkie and Wood, 1991, 1996; Wilkie et al., 1993). The alkalosis-induced disruption to ammonia excretion is likely further aggravated by increased ambient temperature typical of aquatic systems after the wildfire denudes the overhead shading vegetation in riparian habitats (Warren et al., 2022), and this trend is observed in the majority of studies (Paul et al., 2022). Warmer temperature would inevitably elevate basal metabolic demands, increases organismal metabolism, promote faster ammonia production, and decrease the energy budget available for mitigating the IOA-B disturbance (Gomez Isaza et al., 2022). As such, post wildfire ash-input poses a significant IOA-B challenge in need of greater research and consideration. To the best of our knowledge, there have been no studies examining teleost IOA-B response to ash-induced environmental alkalosis, nor their concurrent response to warmer conditions.

The objective of this study was to quantify how ash input can greatly and rapidly induce an acid-base challenge for aquatic organisms, and the fish’s initial physiological response to environmental alkalinization. Our first objective was to identify the amount of ash-input relevant for environmental comparison. We accomplished this by characterizing the water quality parameters of our experimental water and its response to different ash concentration, and determined 0.25% (w/v) as the appropriate ash concentration to use for our study (see below). Our second objective was to determine the biological response to post-wildfire ash-input. Chinook salmon were acclimated to 15 and 20°C to represent temperature downstream or at the site of the burn, respectively. Two weeks later, Chinook salmon response to no ash exposure or after 1, 12, or 24 hours of ash (0.25% w/v) by measuring a suite of blood ionic and acid-base parameters to determine their initial and short-term response to the ash-induced environmental alkalosis.

## Methods

### Fish Husbandry Condition

This experiment was conducted in April and May 2023 in accordance to the protocol no. 23316 in compliance with the Institutional Animal Care and Use Committee (IACUC) at the Center for Aquatic Biology and Aquaculture (CABA) at University of California Davis (UCD). Fall-run Chinook Salmon hatched on December 4^th^, 2021 at the Feather River Hatchery (Oroville, CA, United States) were transferred to the CABA on January 31^st^, 2022, and they were reared in a flow-through well-water system at 13-15°C and fed at 4% body mass per day. Fish (fork length: 17.5 ± 0.2 cm, body mass: 61.0 ± 2.2 g) were acclimated to 15°C or 20°C for at least 2 weeks before experimentation, which took place between March to May, 2023. Fish were starved for 24 hours prior to experimentation.

### Water Quality of Experimental Condition

Experimental water temperature, DO, pH, and salinity were measured daily using YSI 556 MPS (Yellow Springs, Ohio, USA). Temperature was also recorded with temperature loggers at 15-min intervals (Onset Corporation, Cape Cod, MA, USA). Discrete water samples were taken for alkalinity (HACH digital titrator, HACH; Loveland, CO, USA), and an end point was detected using a pH microelectrode (HI1083B, Hanna Instruments, Woonsocket, RI, United States) and meter (HI8424, Hanna Instruments). Turbidity was measured (HACH 2100Q Handheld Turbidity Meter) following manufacturer instructions. Alkalinity, pH, salinity, and temperature values were used to calculate *p*CO_2_ using CO2SYS (version 1.05; Lewis and Wallace, 1998). In addition, experimental water was collected before ash-dosing and after the exposure duration for elemental analysis. Experimental water was filtered (0.45 µm) and stored in a sterile plastic container, mixed at a 13:1 ratio with 1% nitric acid (Certified ACS Plus), stored at room temperature, then later analyzed by the Interdisciplinary Center for Plasma Mass Spectrometry at the University of California at Davis (ICPMS.UCDavis.edu) using an Agilent 8900 ICP-MS Triple Quad instrument (Agilent Technologies, Santa Clara, CA 95051).

### Water Quality of Field Condition

Spot measurements of several aquatic systems local to Davis (California, United States) were used to determine the relevant experimental treatment (explained below). These include measurements collected from fresh rainwater (collected in plastic buckets and measured in the rain) and mud puddles around CABA, river water at the nearby Putah Creek, and lake water at Lake Berryessa (collected at UC Davis Putah Creek Facility) taken between February to April 2023.

### Assessing Ash Impact on Well and Deionized water

Leachate tests were performed on well and deionized (DI) water to illustrate the relationship of ash and alkalinity. Briefly, sieved ash (pore size = 0.841 mm) were mixed with CABA well water (0, 0.1, 0.25, 0.5, 1, 3%) or DI water (0, 0.25%) and stirred with a magnetic stirrer for 5 min. Next, their temperature, DO, pH, salinity, alkalinity, and turbidity were measured as previously described. As a reference, leachate methods detailed in USGS Field sampling guide and (Burton et al., 2016) utilizes a 5% (w/v) mixture (1 g in 20 mL).

### Experimental Condition

The ash used in this experiment were derived from a combination of oak trees burnt in a furnace and local control burning of pomegranate, oak, and redwood trees. An experimental exposure of 0.25 % ash was achieved by mixing 250 g of ash into the 100 L tank. Each tank was stocked at a density of 6-8 fish per tank. To minimize disturbance, ash was mixed with water in an external bucket, then transferred into the experimental chambers using aquarium submersible pumps. CABA well water inflow was halted during the experimental ash exposure. Air bubbling and a second submersible pump within the experimental tanks assisted with water mixing. Ash addition increased the pH of 15 and 20°C treatment water from 8.13 and 8.06 to 9.27 and 9.17, respectively (Table 1). Temperature remained relatively consistent throughout exposure, though the 15°C treatment warmed slightly over time due to the warmer air temperature (Table 1). Ash input increased pH, alkalinity, and salinity, but decreased *p*CO_2_ (Table 1). Sample size at each ash exposure timepoint are as followed: 0 hour (15°C: n=9; 20°C: n=10), 1 hour (15°C: n=9; 20°C: n=5), 12 hours (15°C: n=9; 20°C: n=11), 24 hours (15°C: n=12; 20°C: n=14).

**Table 1:** Water quality of experimental tanks at pre- and post-ash exposure held at 15 or 20°C. Values are mean ± SEM.

|  | 15°C |  | 20°C |  |
| --- | --- | --- | --- | --- |
|  | Pre-Ash Exposure | Post-Ash Exposure | Pre-Ash Exposure | Post-Ash Exposure |
| **pH** | 8.13 ± 0.01 | 9.27 ± 0.09 | 8.06 ± 0.01 | 9.17 ± 0.09 |
| **DO (mg/L)** | 9.98 ± 0.15 | 9.89 ± 0.10 | 8.76 ± 0.15 | 8.00 ± 0.13 |
| **Temperature (°C)** | 15.6 ± 0.1 | 16.5 ± 0.3 | 20.0 ± 0.2 | 19.9 ± 0.1 |
| **Alkalinity ($\mu\text{mol/kg}$)** | 3,954 ± 157 | 4,694 ± 342 | 3,787 ± 245 | 4,732 ± 394 |
| **$p\text{CO}_2$ ($\mu\text{atm}$)** | 1,378 ± 65 | 84 ± 22 | 1,680 ± 114 | 95 ± 29 |
| **Salinity (ppt)** | 0.44 ± 0.002 | 0.54 ± 0.015 | 0.45 ± 0.001 | 0.54 ± 0.009 |

### Blood Sampling and Analysis

We sampled blood without the use of cannulation via a gill irrigation technique described in past studies (Harter et al., 2021; Kwan and Tresguerres, 2022; Davison et al., 2023). After ash exposure reached the designated timepoint, fish were anesthetized in their treatment tank by pouring a benzocaine stock solution through an extended tube that is out of view of the fish to minimize disturbance to achieve a concentration of 75 mg/L benzocaine in the tank. After loss of equilibrium (∼3 min), fish were moved to a surgery table where their gills were irrigated with aerated treatment water with maintenance anesthetic (benzocaine, 30 mg/L). Blood was drawn from a caudal vessel using a heparinized syringe (21 gage needle; 100 IU lithium heparin), placed on ice, then processed within 5 minutes of sampling. All sampling took place between the hours of 8:00 and 13:00.

Whole blood pH (pH_e_) was first measured with a micro pH electrode (HI1083B, Hanna Instruments), then a subset (65 μL; 1-3 sample per fish) was analyzed using the ABL90 Flex Plus (Radiometer, Copenhagen, Denmark) to measure blood *p*CO_2_ and the concentration of Na^+^, K^+^, Cl^−^, Ca^2+^, glucose and lactate. The remainder of the samples were spun for 2 min on a tabletop centrifuge, and the separated red blood cell (RBC) and plasma fractions were flash frozen with liquid N_2_ for later intracellular pH (pH_i_), total ammonia and CO_2_ measurements. RBC pH_i_ was measured using the freeze-thaw technique (Zeidler and Kim, 1977). Following best practices (Baker et al., 2009), pH_i_ was measured with a pH microelectrode (HI1083B, Hanna Instruments) within 2-weeks of the conclusion of the experiment to limit pH change. *p*CO_2_ values were temperature corrected using the following equation (Siggaard-Andersen, 1974).

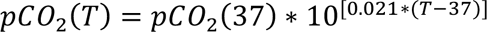

Next, blood pH_e_ and *p*CO_2_ values were used to calculate [HCO_3_^−^] using the Henderson-Hasselbalch equation. The solubility coefficient of CO_2_ (0.0578 mmol l^−1^ Torr ^−1^), ionic strength (0.15 M), and pK_1_ (6.20) were based upon (Boutilier et al., 1984). Plasma total ammonia ([T_Amm_]i.e. [NH_3_ + NH_4_ ^+^]) was determined spectrophotometrically in 25 µL aliquots of flash frozen plasma by enzymatic ammonia assay (Sigma-Aldrich, USA, Catalog number: AA0100). [T_Amm_] was calculated from the delta absorbance at 340 nm wavelength before and after the addition of the enzyme L-glutamate dehydrogenase. Absorbance was measured in Greiner UV-star® 96-well plates using a microplate reader (Infinite® M200 PRO, Tecan, Switzerland). Finally, *p*NH_3_ and [NH_4_^+^] were calculated with the Henderson-Hasselbalch equation using the solubility coefficients and pK_Amm_ values from Cameron and Heisler (1983).

### Statistical Analysis

Statistical analyses were performed using *R* (version 4.0.3) (R Development Core Team, 2013). Water quality parameters were analyzed with two-tailed Student’s t-test, one-way Analysis of Variance (ANOVA), and linear regressions with water source and ash input as covarying factors. Blood and plasma variables were analyzed with Analysis of Covariance (ANCOVA), with temperature and duration of ash exposure as factors. Normality and homogeneity of residuals were assessed through visual inspection of QQ plots and residual boxplots, respectively. An alpha level of 0.05 was used for significance in all statistical tests. Unless noted otherwise, results are reported as mean ± SEM.

## Results

### Part 1: Water Quality Impact from Ash Input

Water quality parameters were measured in stagnant and fresh rain and mud puddles, DI water, CABA well water, Putah Creek, and Lake Berryessa (Supplemental Figure 1). Fresh rainwater (pH ∼6.2, alkalinity ∼40 μmol/kg) is more acidic and less buffered than adjacent mud puddles (pH∼7.3, alkalinity ∼300 μmol/kg), which reflects the slightly alkaline soil reported around Davis, CA (Walkinshaw et al., 2022). DI water (pH ∼7.0, alkalinity ∼40 μmol/kg) is neutral in pH, but has a similar level of alkalinity as rainwater. In contrast, natural overland water bodies such as Putah Creek and Lake Berryessa have pH ∼8.3 and alkalinity ∼1,800 μmol/kg, and these elevated values are likely attributed to mineral absorption from soil and rock erosion over time. Well water has similar pH (∼8.1) to that of overland water, but its alkalinity (∼3,700 μmol/kg) is more than double that of Putah Creek and Lake Berryessa. This is likely because the well water has had the most time to interact with minerals as it permeated through the groundwater system, but *p*CO_2_ levels is elevated because it has not yet equilibrated with the atmosphere giving rise to a similar pH overall. Likewise, freshwater alkalinity, turbidity, and salinity also correlated with time spent interacting in the watershed, though turbidity of DI, Lake Berryessa, and CABA well water were likely low due to removal by filtration systems at UCD.

The relationship between ash input across concentration was examined by mixing CABA well water with 0, 0.1, 0.25, 0.5, 1, and 3% (w/v) locally burned ash. Ash input induces a logarithmic increase in water pH and alkalinity, and a logarithmic decrease in *p*CO_2_ (Figure 1A-C). In contrast, ash input linearly increased salinity and turbidity (Figure 1D, E), and did not affect DO (Figure 1F). The magnitude of pH change induced by ash input was dependent on the starting alkalinity of the water. To demonstrate this, DI and well water response to an ash input of 0.25% (w/v) were compared. DI water increased pH much more (∼7.0 to ∼10.5) than the more buffered well water (∼8.1 to ∼9.2) during ash exposure (Figure 2A). In contrast, the rate of ash-induced changes in alkalinity, *p*CO_2_, salinity, and turbidity were relatively similar (Figure 2B-E). Finally, ash input did not impact DO (Figure 2F).

**Figure 1:**
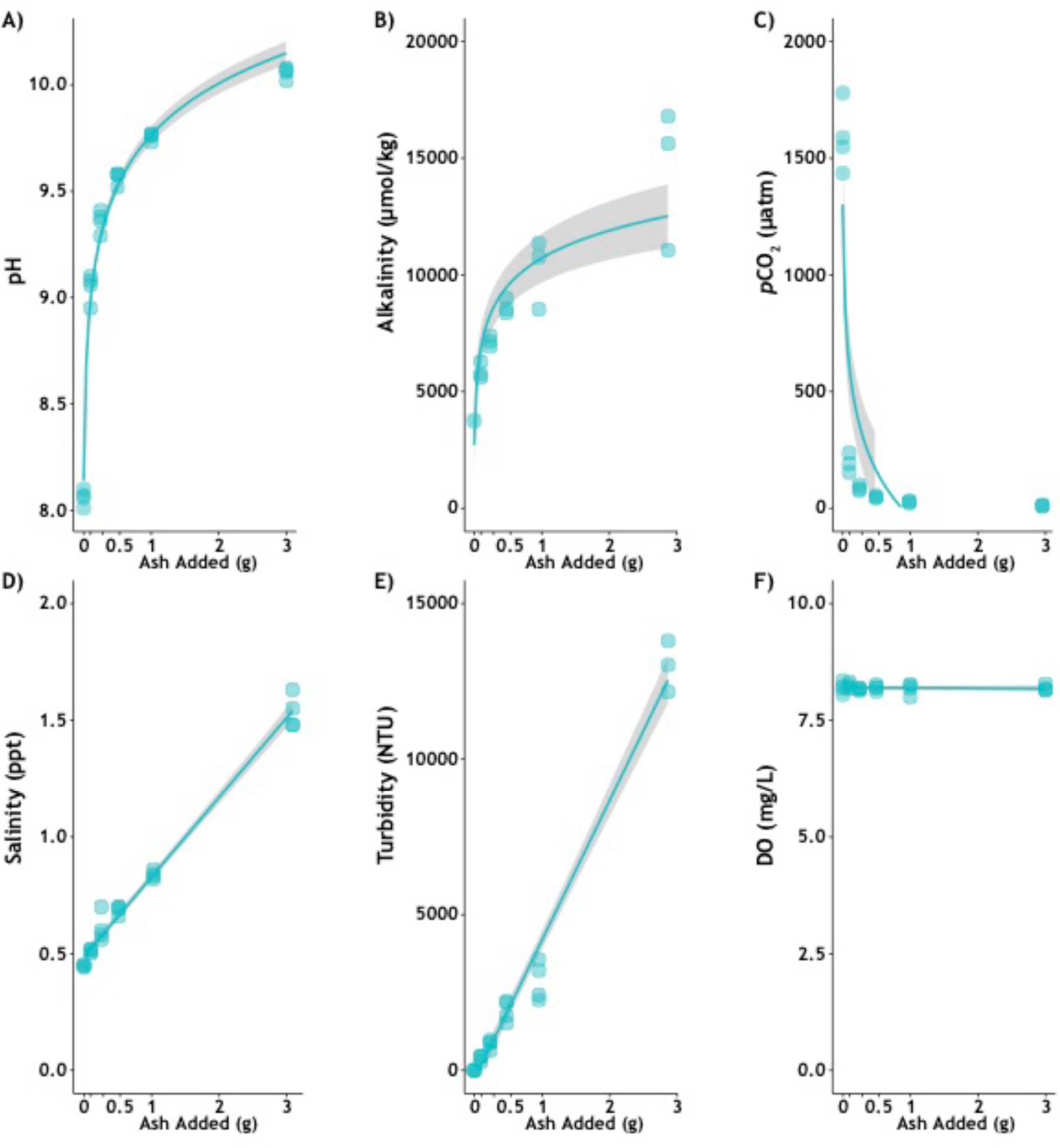
CABA well water and ash dosing curve. The impact of adding 0.1, 0.25, 0.5, 1, and 3% ash (w/v) on CABA well water A) pH, B) alkalinity, C) *p*CO_2_, D) salinity, E) turbidity, and F) dissolved oxygen levels. Shaded area denotes 95% confidence interval.

**Figure 2:**
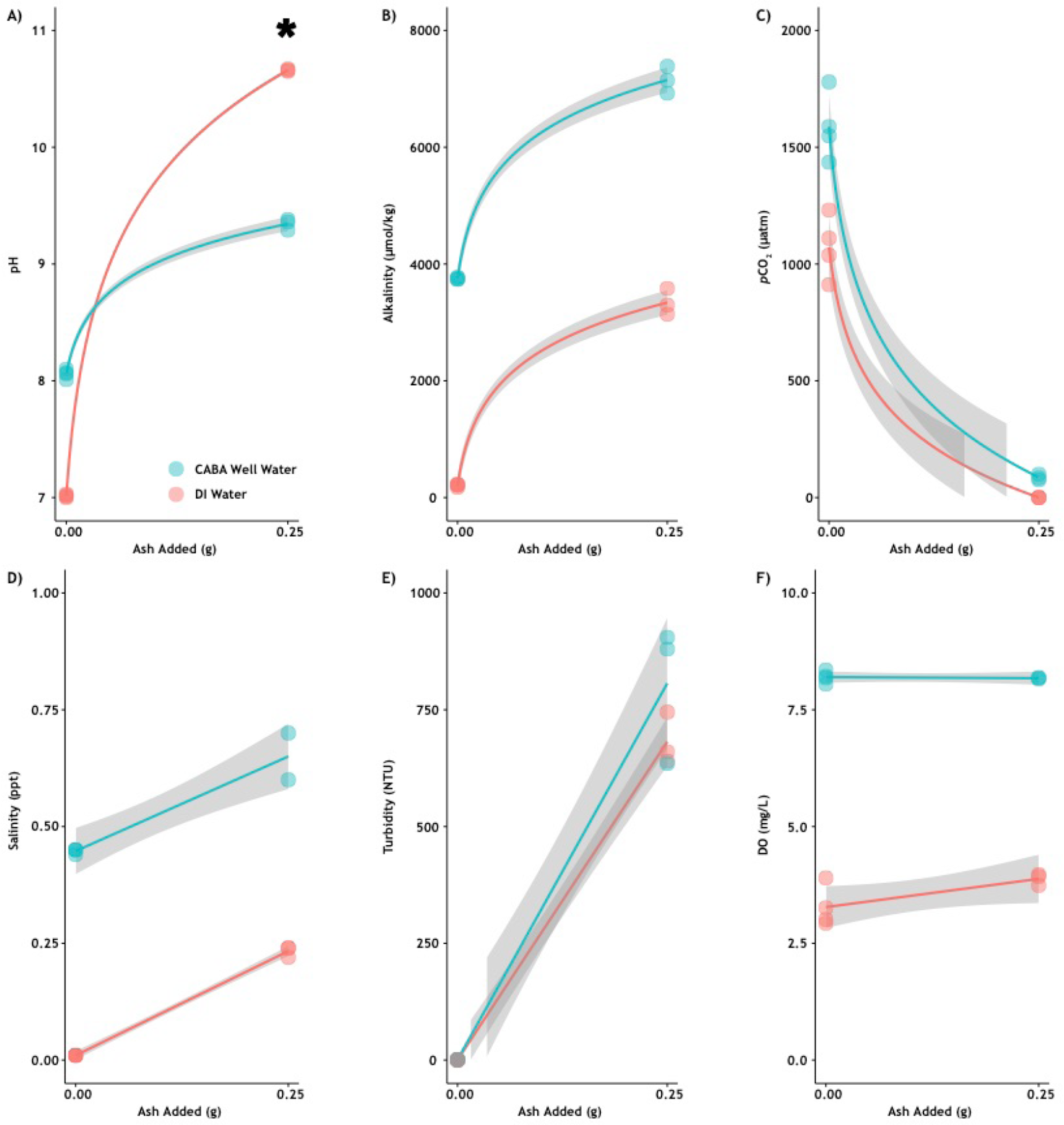
Influence of salinity on ash-induced water chemistry change. Comparison of DI and CABA well water with and without 0.25% (w/v) ash input on A) pH, B) alkalinity, C) *p*CO_2_, D) salinity, E) turbidity, and F) dissolved oxygen levels. Shaded area denotes 95% confidence interval.

ICP-MS analysis identified 25 of 31 elements within detectable range (Supplemental Figure 2-4). None of the 25 elements was affected by temperature, nor were there significant decline in concentration after ash input. Ash input significantly increased the concentration of 13 elements: Al, B, Ba, Cr, K, Li, Mo, Mn, Ni, P, Sb, Se, and Sn (Supplemental Figure 2-4).

### Part 2: Blood Response to Ash Exposure in 15 and 20°C acclimated Fish

Because the majority of past wildfire studies responded with an increase of ∼1 pH units (Paul et al., 2022), we selected the concentration of 0.25% (w/v) was selected as our experimental condition. This value would induce a pH increase from ∼8.0 to ∼9.2, which is slightly higher than the EPA recommended upper limit of ∼9.0 (U.S. Environmental Protection Agency., 2013) yet demonstrated to be within tolerable range by the related rainbow trout (Wilkie and Wood, 1991). Chinook salmon yearlings acclimated to 15 or 20°C were exposed to 0, 1, 12, or 24 hours of 0.25% (w/v) ash exposure. In total, we observed 20% (6 of 30) and 33.3% (10 of 30) mortalities in the 15°C and 20°C treatment, respectively, all of which occurred between 1 to 12 hours of ash exposure.

Prior to ash exposure, the pH_e_ of salmon reared at 15°C (7.81 ± 0.02) were slightly but not significantly lower than those reared at 20°C (7.92 ± 0.02) in fishes (*p* = 0.2148; Figure 3A). In contrast, RBC pH_i_ (15°C: 7.55 ± 0.02, 20°C: 7.56 ± 0.02) were extremely similar (*p* = 1.0000; Figure 3B). Salmon reared at 20°C had significantly higher *p*CO_2_ (4615 ± 269 μatm; *p* = 0.0089) than 15°C acclimated fish (3583 ± 251 μatm; Figure 3C). Warmer temperature also significantly elevated baseline plasma [HCO_3_^−^] (*p* < 0.0001) (15°C: 4.75 ± 0.13 mM; 20°C: 7.06 ± 0.16 mM; Figure 3D).

**Figure 3:**
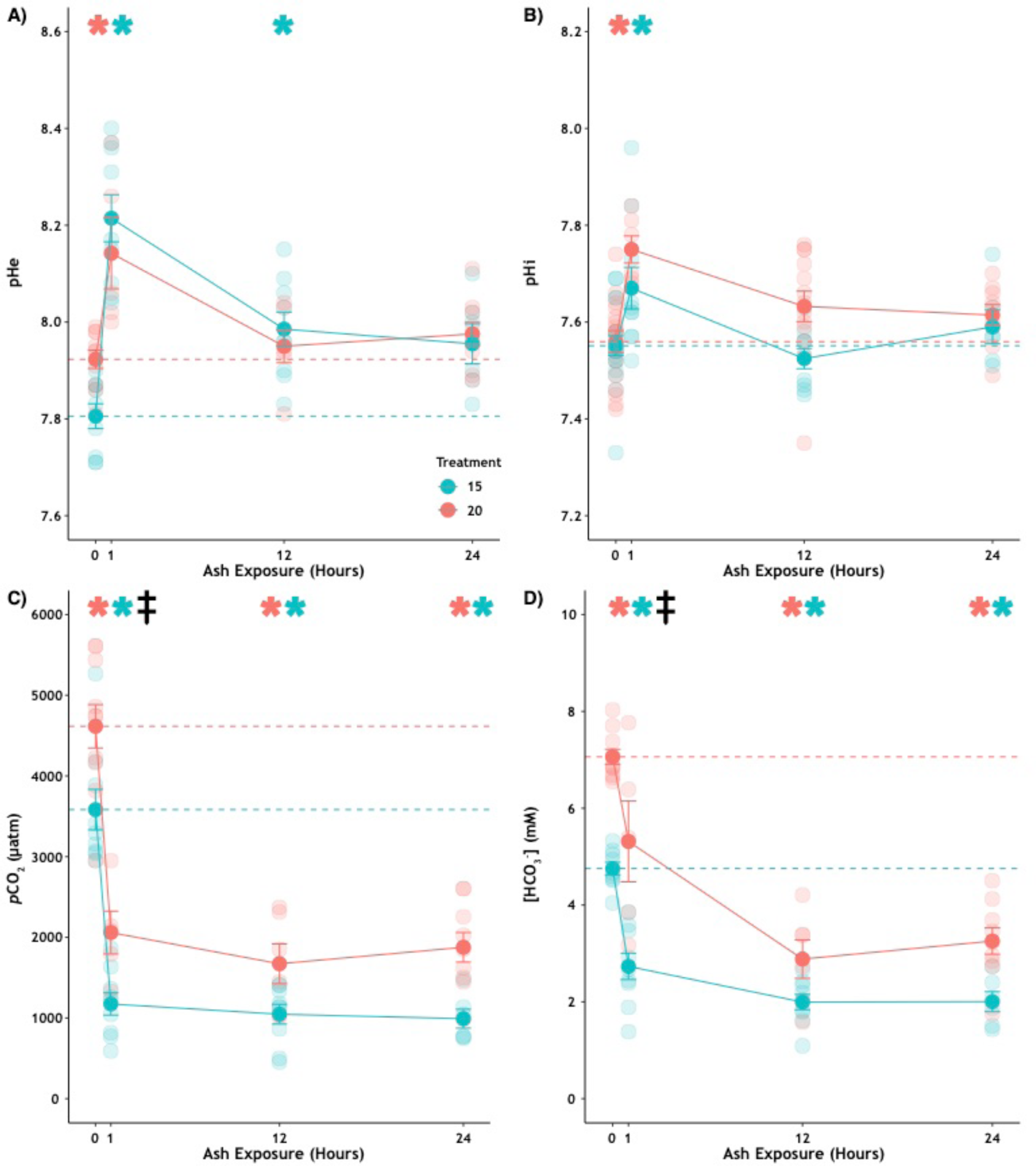
Blood acid-base response to ash-induced pH ∼9.2 exposure at 15 and 20°C. Salmon blood A) pHe, B) pHi, C) pCO_2_ and D) [HCO_3_^−^] over the 0, 1, 12, and 24 hours of ash exposure. Values are mean ± SEM. Asterisk (teal = 15°C, salmon = 20°C) indicates significance (α = 0.05) from respective control (0 hour exposure), which is represented as a dotted line. Black diesis (double dagger) indicates significance between the 15 and 20°C 0-hour controls.

Ash input rapidly increased water pH from ∼8.1 to ∼9.2 and simultaneously decreased *p*CO_2_ from 1,400 – 1,700 to ∼100 μatm (Table 1), challenging the salmon with acute environmental alkalosis. This study shows that 1 hour was not enough time for the salmon to mitigate the acid-base disturbance: both salmon pH_e_ (15°C: *p* < 0.0001, 20°C: *p* = 0.0051) and pH_i_ (15°C: *p* = 0.0386, 20°C: *p* = 0.0057) were significantly elevated compared to their respective baselines (Figure 3A, B). However, salmon acclimated to 20°C returned to baseline pH_e_ levels after 12 hours (12 hours: *p* = 0.9995), whereas salmon acclimated to 15°C needed 24 hours to no longer be significantly different from baseline (24 hours, *p* = 0.1214). Salmon RBC pH_i_ also significantly rose (15°C: *p* = 0.0386, 20°C: *p* = 0.0057), but both treatments had fully recovered by 12 hours of exposure. In contrast, blood *p*CO_2_ (15°C and 20°C: *p* < 0.0001) and [HCO_3_^−^] (15°C: *p* < 0.0001, 20°C: *p* = 0.0050) significantly decreased after 1 hour of ash exposure (Figure 3C, D), and they remained low or further declined with prolonged exposure.

Plasma [Na^+^], [K^+^], and [Ca^2+^] were not significantly affected by ash exposure (Figure 4A, C-D). In contrast, plasma [Cl^−^] was differentially affected by ash exposure (Figure 4B): [Cl^−^] was significantly lower than baseline after 12 hours of exposure in 15°C salmon (*p* = 0.0041), and significantly higher than baseline after 24 hours of exposure in 20°C salmon (*p* = 0.0300). Plasma glucose was significantly increased with ash exposure in 15°C fish after 24 hours of exposure (*p* < 0.0001) and in 20°C salmons after both 12 hours (*p* = 0.0001) and 24 hours of exposure (*p* = 0.0002; Figure 5A). In general, plasma lactate in both temperature treatments spiked at 1 hour (15°C: *p* = 0.0565, 20°C: *p* < 0.0001) and decreased over the next 12 (15°C: *p* = 0.7302, 20°C: *p* = 0.1098) and 24 hours (15°C: p = 0.6876, 20°C: *p* = 0.0167), but only those of 20°C salmon was statistically significant (Figure 5B).

**Figure 4:**
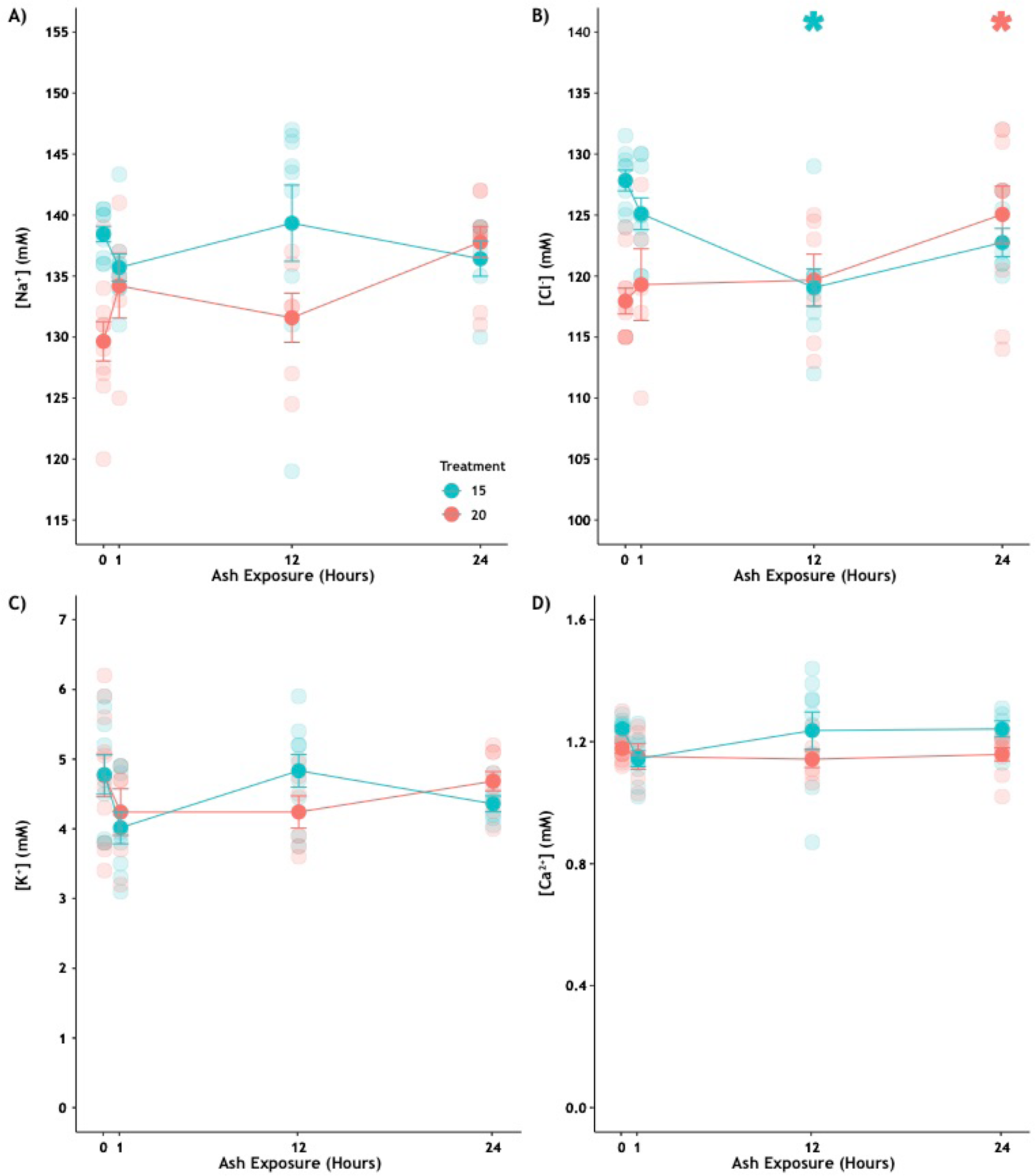
Plasma ion response to ash-induced pH ∼9.2 exposure at 15 and 20°C. Salmon plasma A) [Na^+^], B) [Cl^−^], C) [K^+^] and D) [Ca^2+^] over the 0, 1, 12, and 24 hours of ash exposure. Values are mean ± SEM. Asterisk (teal = 15°C, salmon = 20°C) indicates significance (α = 0.05) from respective control (0 hour exposure).

**Figure 5:**
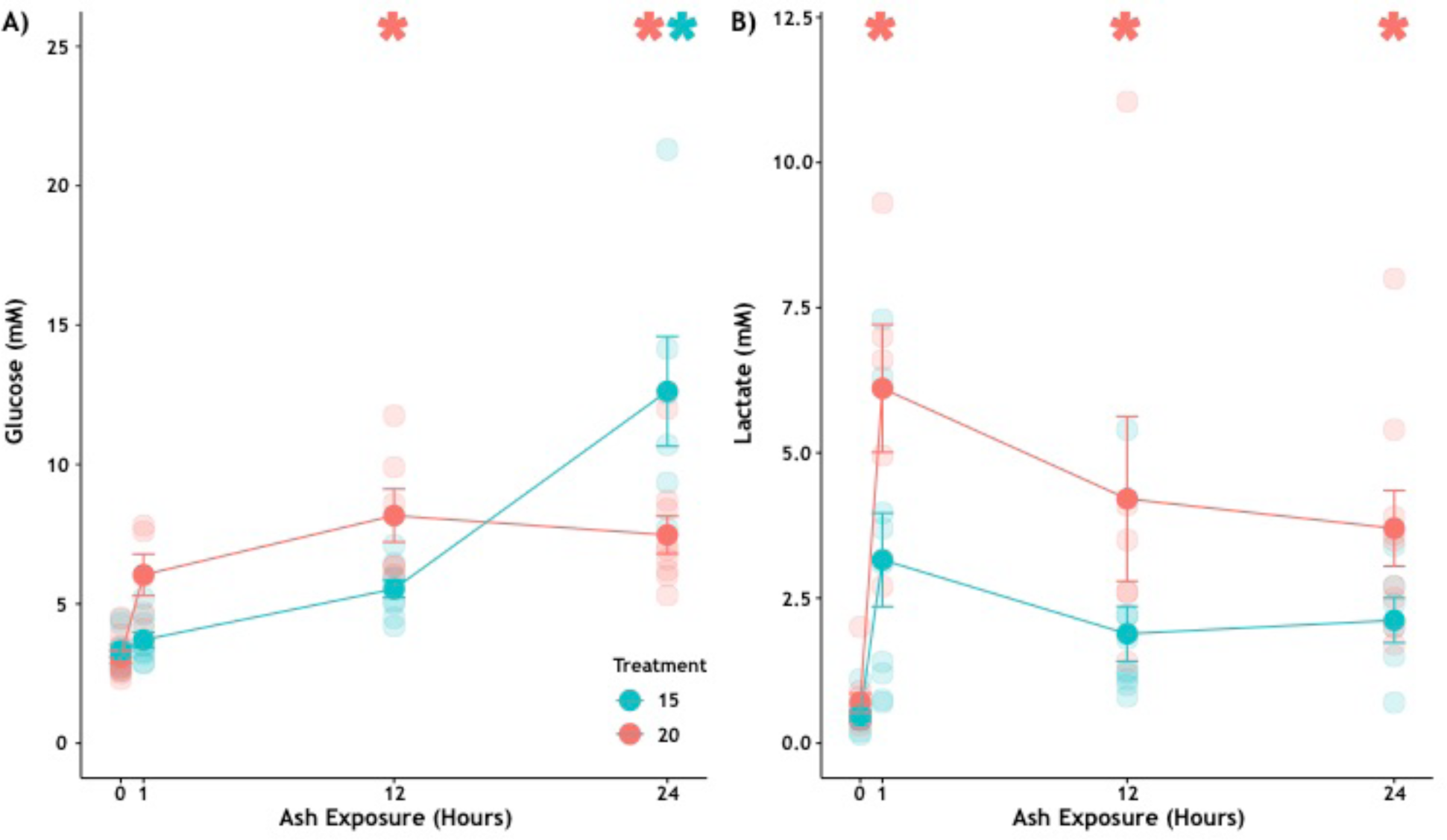
Plasma glucose, lactate, and ammonia response to ash-induced pH ∼9.2 exposure at 15 and 20°C. Salmon plasma A) glucose and B) lactate over the 0, 1, 12, and 24 hours of ash exposure. Values are mean ± SEM. Asterisk (teal = 15°C, salmon = 20°C) indicates significance (α = 0.05) from respective control (0 hour exposure).

Plasma total ammonia generally increased with ash exposure (Figure 6A), but the two temperature treatments exhibited different response patterns: plasma total ammonia in 15°C salmon gradually increased until it was significantly higher at 24 hours of exposure (*p* = 0.0340), whereas 20°C salmon experienced a 7.5-fold increase after 1 hour of exposure (*p* < 0.0001), remained significantly ∼3-fold elevated at 12 hours of exposure (*p* = 0.0105), and finally tapered down to non-significant levels at 24 hours of exposure (Figure 6A). Plasma [NH_4_ ^+^] followed a similar pattern: [NH_4_^+^] in 15°C salmon was significantly elevated after 24 hours of ash exposure (p = 0.030), whereas [NH_4_^+^] in 20°C salmon was significantly elevated after 1 (*p* < 0.0001) and 12 hours of exposure (*p* < 0.0001), but returning back to control levels by 24 hours (*p* = 0.6836; Figure 6B). Finally, [NH_3_] in 20°C salmon was significantly elevated at 1 (*p* < 0.0001) and 12 hours (*p* < 0.0001), but returned to control levels by 24 hours of exposure (Figure 6C). In contrast, [NH_3_] in 15°C salmon was not significantly elevated relative to control levels regardless of ash exposure duration (Figure 6C).

**Figure 6:**
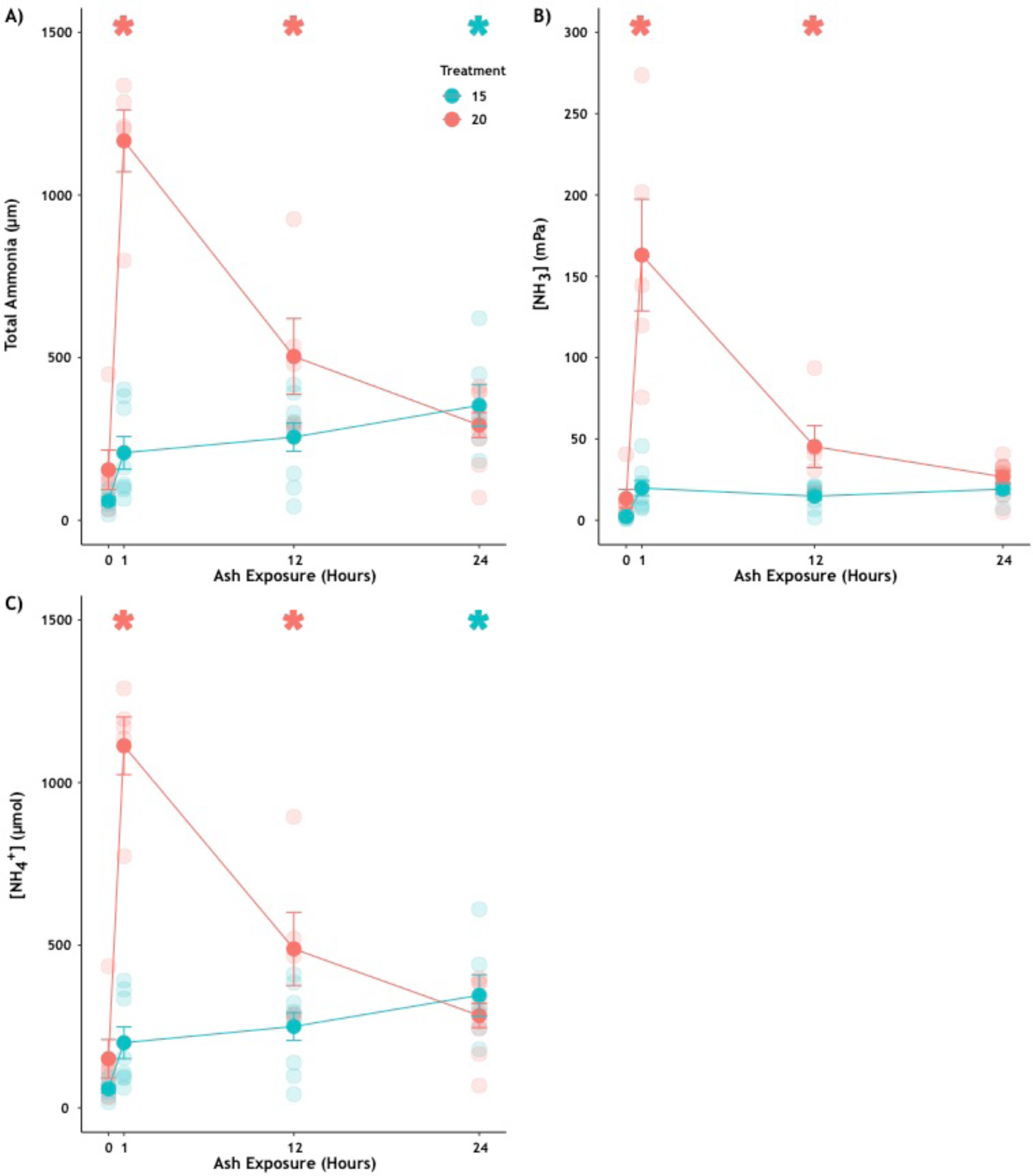
Plasma total ammonia, [NH_3_], and [NH_4_ ^+^] response to ash-induced pH ∼9.2 exposure at 15 and 20°C. Salmon plasma A) total ammonia, B) [NH_3_], and C) [NH_4_^+^] over the 0, 1, 12, and 24 hours of ash exposure. Values are mean ± SEM. Asterisk (teal = 15°C, salmon = 20°C) indicates significance (α = 0.05) from respective control (0 hour exposure).

## Discussion

In agreement with the classic Davenport acid-base physiology and Wilkie & Wood (1991), exposure to pH ∼9.3 initially induced an elevation in blood pH_e_ and pH_i_, and a simultaneous reduction in plasma *p*CO_2_ and [HCO_3_^−^]. Most salmon were able to recover their pH_e_ and pH_i_ for the ash-induced environmental alkalinization after 12-24 hours of exposure, which matches the response time of the rainbow trout (Wilkie and Wood, 1991). One potential mechanism for pH recovery is through the coordinated effort of apical anion exchanger (AE), cytoplasmic carbonic anhydrase, and basolateral Na^+^/H^+^ Exchanger (NHE1) and electrochemically driven by basolateral vacuolar-type H^+^-ATPase similar to base-secreting gill ionocytes in elasmobranchs (Tresguerres et al., 2005; Roa et al., 2014) or basolateral NKA as in teleost intestinal epithelium (Grosell and Genz, 2006). In concept, the increased ions leached from the ash should ease IOA-B recovery. Yet not all of the salmon were successful in this endeavor: we observed a 20% and 33.3% mortality rate in Chinook salmon reared at 15°C and 20°C, respectively. The timing of the mortality occurred between 1 and 12 hours of exposure, and the greater mortality at the warmer temperature cannot be explained by worse acid-base disturbance (in fact it was less disturbed in the warmer fish). Instead, the greater mortality may correlate better with the rapid and substantial increase in plasma total ammonia. Although not directly comparable due to differences in experimental parameter and species, mortality appeared to be greater when water alkalization was induced with ash input rather than NaOH (Wilkie and Wood, 1991) or using naturally occurring alkaline lake water (Wilkie et al., 1994); Wilkie and Wood (1991) reported one rainbow trout perished during 72 hours of exposure to pH 9.5, and in a subsequent experiment reported in the same study no rainbow trout mortality during a 5-week exposure to pH 9.5. Despite this, Wilkie and Wood (1991) concluded that high pH exposure does “render the fish more susceptible to other stresses” as they observed recovery from surgical procedures (e.g. cannulation) was more successful when kept at a lower pH (8.15).

Salmon in the 15°C treatment appeared to have experienced greater alkalosis, and this may explain why salmon reared at 20°C appear to have recovered their pH_e_ more quickly and completely than those reared at 15°C. One possibility is that salmon reared at 20°C are generating CO_2_ at a faster rate and therefore potentially have access to more H^+^ to recover pH_e_ more quickly (providing they can excrete the HCO_3_^−^ generated by CO_2_ hydration). Interestingly, past results showed rainbow trout reared at 15°C (the colder temperature we used) and challenged with environmental alkalinization (pH 9.5) also could not return to their baseline pH_e_ of ∼7.83, and instead stabilized their pH_e_ at 7.97 (Wilkie and Wood, 1991). These results suggest that although warmer temperature may increase mortality, their higher metabolic rate (and by extension basal *p*CO_2_) could facilitate pH recovery by promoting H^+^ synthesis.

Recovery from environmental alkalosis requires the excretion of plasma HCO_3_^−^ and retention of H^+^ to return blood pH to nominal levels. Past studies typically find fishes exposed to environmental alkalinization between 10°C to 15°C have decreased [Na^+^] and [Cl^−^] as well as elevated [K^+^] by 8 hours of exposure, with these effects persisting up to 72 hours of exposure (Wilkie and Wood, 1991; Hemming and Hanson, 1992; Wilkie et al., 1993; Scott et al., 2005). In contrast, the present study finds Chinook salmon plasma ion responses to be relatively muted, with only [Cl^−^] exhibiting a divergent response: salmon reared at 15°C had significantly less [Cl^−^] by 12 hours of ash exposure, whereas salmon reared at 20°C had significantly greater [Cl^−^] by 24 hours of ash exposure. Differences in experimental design may explain the muted ionic responses: ash-input increases the ions available for IOA-B regulation (e.g. ∼8,000-fold increase in environmental K^+^ levels; Supplemental Figure Elemental 3), and warmer acclimation temperature (and by extension greater metabolic rate) leads to higher basal *p*CO_2_ level and greater metabolic H^+^ generation. Future studies should continue to explore the interactions between acid-base regulation in an environment with greater ion availability.

Acclimation temperature appears to have influenced plasma total ammonia, [NH_3_], and [NH_4_^+^] levels throughout ash exposure. Chinook salmon in the 15°C treatment experienced a gradual accumulation of ammonia, with levels that were not significantly different until 24-hours of exposure. This gradual rise over 24 hours was similar to previous alkaline water exposure studies at somewhat higher water pH levels (e.g. Wilkie and Wood, 1991; Wilkie et al., 1994; McGeer and Eddy, 1998). In contrast, Chinook salmon in the 20°C treatment experienced a ∼7.5-fold spike at 1 hour of exposure to ∼1200 µM total ammonia, which remained highly elevated at 12 hours, and returned to baseline levels after 24 hours of exposure. This may be linked to the higher observed mortality in the 20°C salmon as the inability to regulate ammonia during high pH exposure has been attributed to fish mortality in past studies (Wilkie et al., 1993; Wilkie and Wood, 1996). Interestingly, the peak level reached here (1200 µM in just 1 hour) was substantially higher and reached far faster than in those previous studies (250-600 µM) in which the water pH levels used were somewhat higher than in the present ash-exposure study (pH 9.4-10.5; Wilkie and Wood, 1991; Wilkie et al, 1994; McGeer and Eddy, 1998). As such, while higher temperature may assist in pH recovery (see above), it also may lead to a greater ammonia challenge during high pH exposure. It additionally suggests that ash may cause such physiological impacts in a way that cannot be explained by high water pH alone. This may relate to the complex mixture of metals and organic compounds also released into freshwater from ash, but the precise mechanisms and causative agents are beyond the scope of the present study (although see below).

To rapidly address the acute acid-base challenge, salmon appeared to have released glucose from glycogen stores and upregulated anaerobic respiration. However, the strategy appears to be temperature specific: salmon in the 20°C treatment have a lower aerobic scope than their 15°C counterparts (Zillig et al., 2023), so they may have 1) hastened their release of glucose from glycogen stores and 2) upregulated metabolic proton production (as evident by plasma lactate build-up (Robergs et al., 2004) to aid in rapid recovery of pH_e_. In contrast, salmon in the 15°C treatment did not significantly upregulate their glucose until 24 hours of exposure, and lactate levels were consistently about half those in fish at the warmer temperature. Future studies are needed to determine whether the metabolic protons were upregulated to assist with pHe recovery. Moreover, respirometry studies are necessary to determine whether the salmon’s temperature-dependent aerobic capacity influence their pHe recovery, and whether the slowing of ventilation rates could have accumulated greater CO_2_ in an effort to acidify their blood as our current methods using artificial gill ventilation necessary to collect blood samples for physiologically-relevant IOA-B values could have masked potential blood *p*CO_2_ impact.

In the wild, fishes (especially those trapped in lakes and reservoirs) would likely experience ash exposure at longer duration than those in the present study. Despite surviving the initial environmental alkalinization, the potential impacts of trace metal accumulation and their putative inhibition of IOA-B regulation could have long-lasting impacts on the fish (reviewed in Wood et al., 2012a, b). For instance, Cr exposure has been shown to induce oxidative stress and negatively impact DNA integrity in the gill and kidney of the European eel (*Anguilla anguilla L.*) (Ahmad et al., 2006). In addition, weeks to month-long exposure to Cu have been shown to induce apoptosis of gill ionocytes and overall lower gill NKA activity (Li et al., 1998; Lorin-Nebel et al., 2013), whereas weeks-long exposure to Li has been shown to decrease fat stores, but did not affect gill NKA activity (Tkatcheva et al., 2007). Even if the concentration for each element is sublethal, the additive effect along with other stressors could accumulate to induce greater impacts or even mortality. Moreover, the elemental signatures and their concentrations should greatly vary depending on factors including (but not limited to) wildfire intensity, vegetation, soil type, anthropogenic input (e.g. fire retardants, buildings, vehicles) – and thus warrant species- and region-specific examination. Behavioral response such as boldness and shoaling have also been shown to be affected by wildfire-ash exposure (Gonino et al., 2019), and downstream food web impacts and ecological dynamics (Spencer et al., 2003) should also be investigated.

### Environmental Relevance and Potential Management Strategies

Over the past decades, numerous studies have examined fish responses to high-CO_2_ low-pH conditions in the context of ocean acidification and aquaculture (Ellis et al., 2016; Tresguerres and Hamilton, 2017). Perhaps due to a lack of recognized environmental relevance, responses of aquatic organisms to low-CO_2_ high-pH conditions remain relatively unexplored, leaving an abundance of exciting research questions that need answering. Would the absence of plasma accessible carbonic anhydrase (which could reduce the magnitude of blood pH_e_ increase) in fish such as sturgeon be more tolerant of environmental alkalosis? This could potentially explain their survival in the Klamath River, which naturally reaches pH ∼10 during the summer (U.S. Geological Survey, 2016). Moreover, what are some conservation management solutions that could reduce the impacts of post-wildfire ash-input in areas inhabited by endangered fishes and other aquatic organisms? Undoubtedly, further comparative physiology and conservation biology research is necessary as wildfires become increasingly common due to global climate change.

The present study uses high alkalinity well water, which would dampen ash-induced pH alkalinization as well as provides more counterions for IOA-B regulation. Therefore, our results may be a conservative estimate of water quality and fish responses to ash input. Many natural systems, especially those found at mid- to high-elevation and sourced with snowmelt, have very low alkalinity levels (Stoddard, 1987; Eilers et al., 1990; Catalan and Camarero, 1993; Clow et al., 1996) and are experiencing more rapid warming than lower elevation aquatic systems (Zhi et al., 2020). Thus, these systems are potentially more susceptible to post-wildfire ash-induced pH changes, which could be detrimental to fishes that are endemic to remote mid- to high-elevation stream habitats like the Paiute Cutthroat Trout (*Oncorhynchus clarkia seleniris*), the California state fish golden trout (*Oncorhynchus aguabonita*), and various salmon runs returning from the ocean. Moreover, and as mentioned earlier in this study, organisms living in systems with naturally high pH levels (e.g. green sturgeon in Klamath River, Lahontan cutthroat trout in Pyramid Lake) and migratory species with rigid reproductive strategies (e.g. salmon) are also more susceptible to the alkalinization-linked impacts of post-wildfire ash-input. Besides baseline water pH and alkalinity, there are many other variables that could influence post-wildfire pH responses including wildfire intensity (Santín et al., 2015; Sánchez-García et al., 2023), including soil pH and composition (Marcotte et al., 2022), amount of rainfall or snowmelt (Rhoades et al., 2011), watershed size and slope (Neary et al., 2003), and algal biomass (Hohner et al., 2019). Taken together, the impacts of wildfire on aquatic watersheds will likely be system-specific, and subsequent organismal response species-specific.

Vulnerable habitats can be identified with greater pH and alkalinity monitoring effort. Monitoring stations equipped and maintained with calibrated pH probes would provide valuable information on pre-wildfire baseline and natural pH variation, which would in turn help determine whether further management action is necessary following a wildfire. For instance, the release of dam water could help dilute the ash-induced alkalization, as well as to mitigate potential co-stressors such as high temperature, low oxygen, and the accumulation of heavy metals. Moreover, the opportunistic capture and removal of migratory species (e.g. returning adults, fertilized redds) from areas of concerns to rear in another area or aquacultures could be another method to safeguard the population. These two strategies are already employed for other scenarios, and with greater monitoring could be applied to help mitigate exposure to excessive ash-input. In principle, ash could be neutralized to prevent the system from reaching lethal pH levels to protect endangered and endemic species. However, many of these strategies may only be employable in the aquaculture setting and/or require additional investigation before they can be safely deployed. For instance, the addition of H^+^ through an acidifying agent (e.g. HCl, CaSO_4_, CH_3_COOH) should lower water pH levels; however, past experiments with freshwater aquaculture ponds have revealed direct acidification is too expensive or temporary to be of practical use (Pote et al., 1990; Tucker and D’Abramo, 2008). One current best practice to reduce high pH in freshwater aquaculture includes the addition of cracked corn or soybean meal, of which their decay would generate CO_2_ through microbial activity (Pote et al., 1990; Tucker and D’Abramo, 2008). However, the decay of organic matter would also deplete O_2_ – which could be mitigated in aquaculture settings with air bubblers but is not feasible for the natural environment. Another technique proposed for freshwater aquaculture is to acidify high pH water by adding aluminum sulfate (Al_2_(SO_4_)_3_), which not only produces H^+^ but also coalesces algae and suspended particles (Tucker and D’Abramo, 2008; Hohner et al., 2019). However, whether aluminum sulfate could neutralize and/or reduce suspended ash in natural riverine systems is not known. Moreover, aluminum is toxic to organisms (especially in freshwater habitats) and can inhibit both active ion-uptake and accelerate passive ion-losses (reviewed in Wilson, 2012). As such, the use of aluminum sulfate must be critically examined and its downstream impacts and interaction with other chemicals present in the natural environment must be robustly explored before being considered as a management tool. Altogether, there is a great need to further investigate aquatic organismal responses to post-wildfire, and to synthesize and develop relevant management strategies to help them survive in an increasingly wildfire-prone climate.

## Acknowledgement

The authors thank Dennis Cocherell (UC Davis) and Linda Deanovic (UC Davis) for coordinating and supplying the ash used in this experiment, and we are grateful to Levi Lewis (UC Davis) for supplying elemental analysis reagents. Elemental analysis was performed at the UC Davis Interdisciplinary Center for Plasma Mass Spectrometry, a Campus Research Core Facility, using an Agilent 8900 ICP-MS Triple Quad purchased with funding from the UC Davis Research Core Facilities Program’s Campus Research Core Facility Enhancement Funding Program managed by the UC Davis Office of Research. Plasma total ammonia analyses were performed at the University of Exeter funded by a BBSRC grant (BB/W018039/1) to RWW. We also thank Brendan Lehman from NOAA Southwest Fisheries Science Center in Santa Cruz for providing manuscript feedback and environmental context to our post-wildfire ash input scenario. GTK was funded by the Delta Stewardship Council Delta Science Program under Grant No. (21045), and NAF was funded by University of California, Davis Agricultural Experiment Station (grant #2098-H). The contents of this material do not necessarily reflect the views and policies of the Delta Stewardship Council, nor does mention of trade names or commercial products constitute endorsement or recommendation for use.

## Data Availability

The data that support the findings of this study are openly available in Dryad (DOI: TBD).

