## Supplemental Material for "Fish Blood Response to Ash-Induced Environmental Alkalinization, and their Implications to Wildfire-Scarred Watersheds"

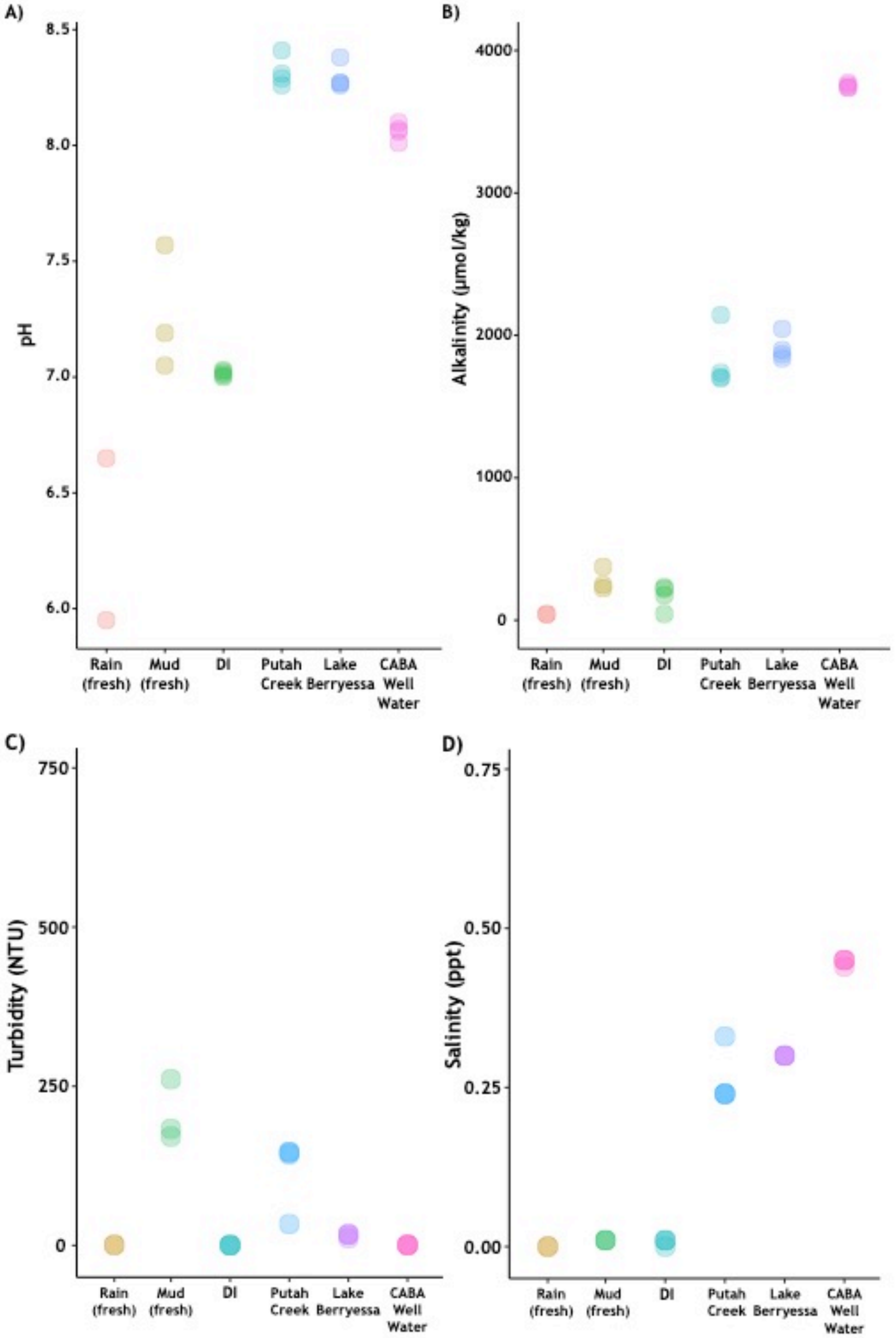


**Supplemental Figure 1:** Spot pH and alkalinity measurements in select water bodies in February and March 2023. A) pH, B) alkalinity, C) turbidity, and D) salinity measurements from rain (stagnant and fresh), mud puddles (stagnant and fresh), DI water, Putah Creek, Lake Berryessa, and well water from at the Center for Aquatic Biology and Aquaculture, University of California Davis.

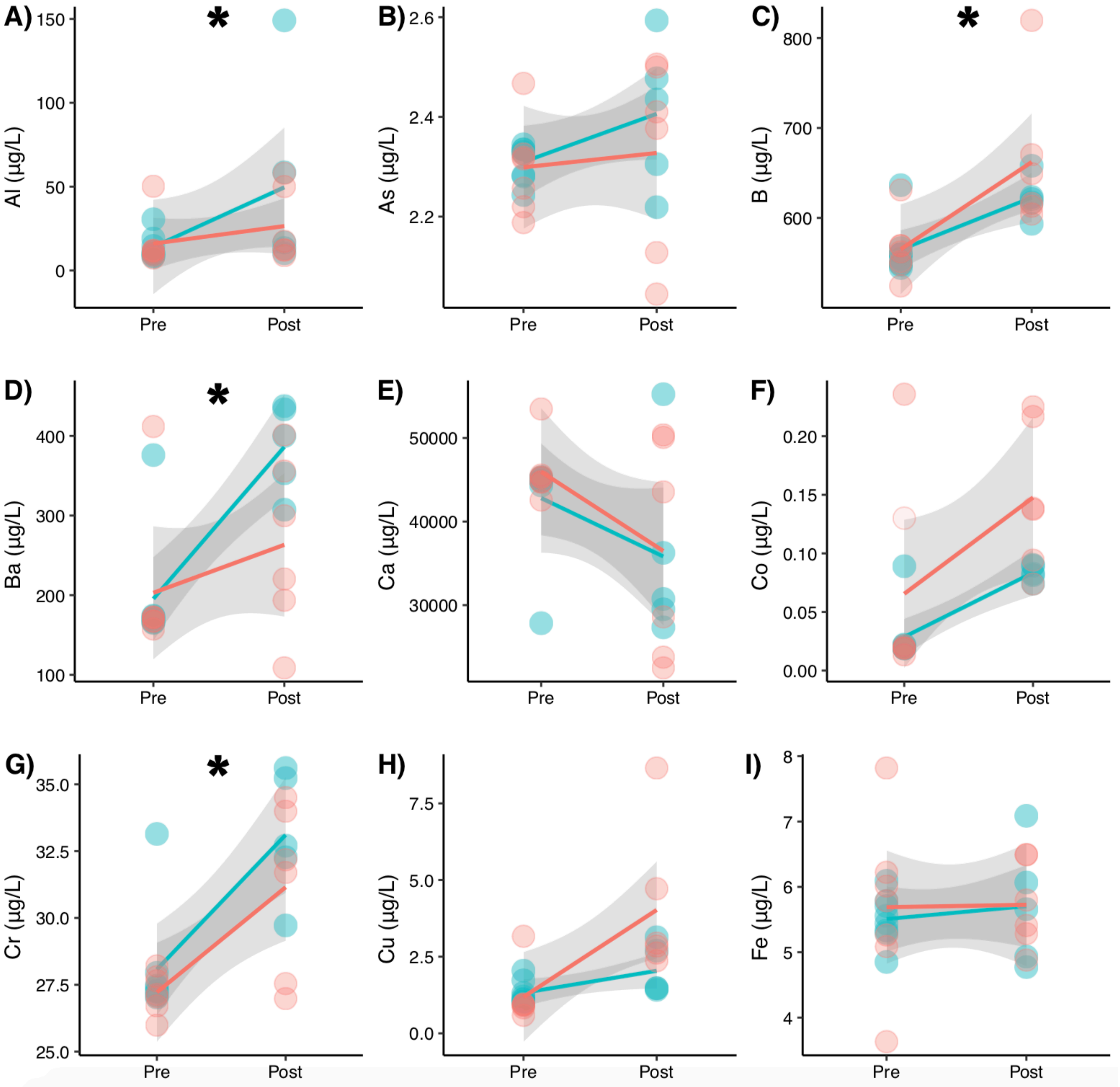


**Supplemental Figure 2:** Change in elemental concentration from ash input. Change in A) Al, B) As, C) B, D) BA, E) Ca, F) Co, G) Cr, H) Cu, I) Fe concentration after ash input into experimental treatment water. No temperature difference (teal = 15°C, salmon = 20°C) was detected. Black asterisk indicates significance (α = 0.05) between pre- and post-ash input. Shaded area denotes 95% confidence interval.

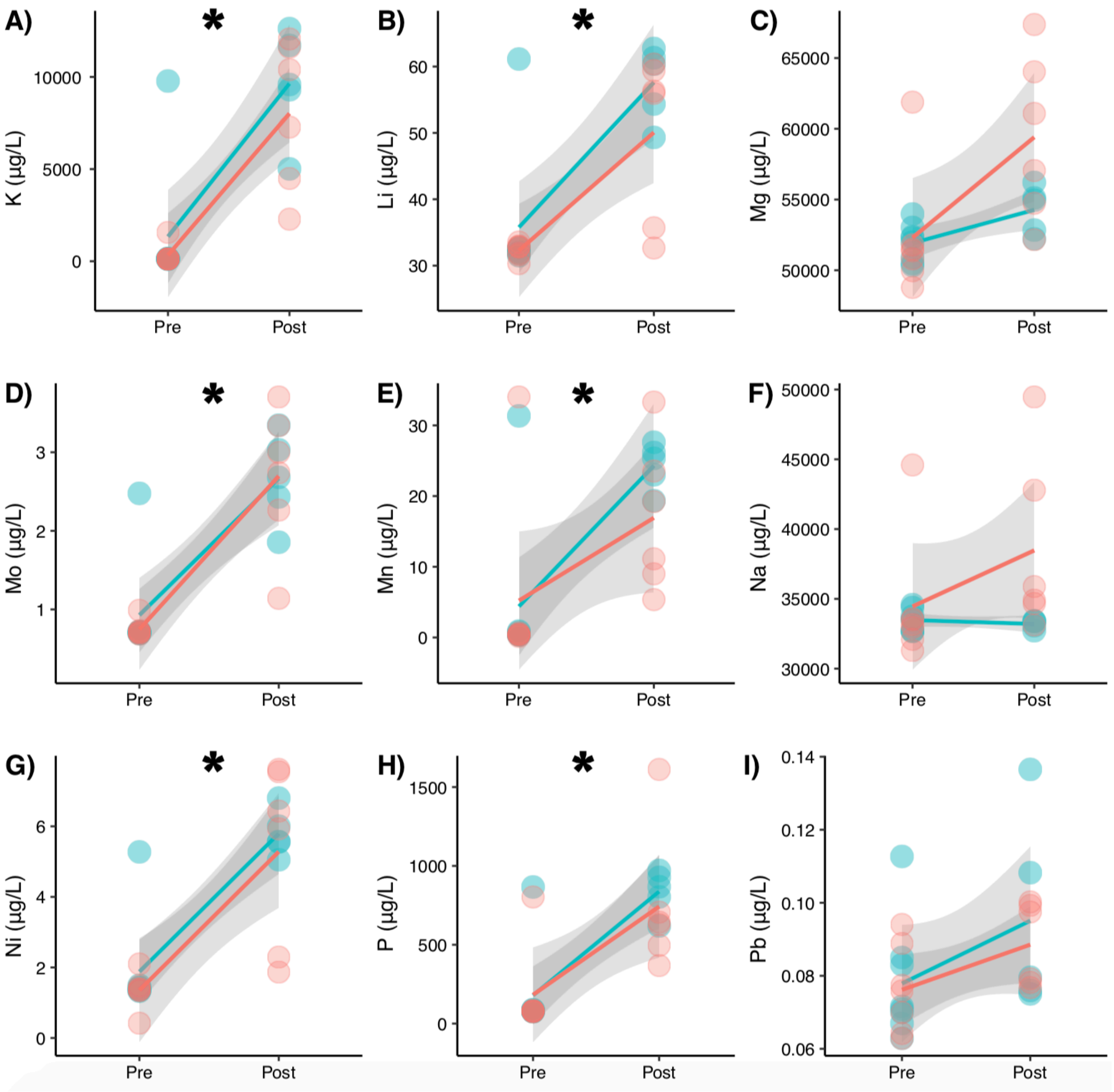


**Supplemental Figure 3:** Change in elemental concentration from ash input. Change in A) K, B) Li, C) Mg, D) Mo, E) Mn, F) Na, G) Ni, H) P, I) PB concentration after ash input into experimental treatment water. No temperature difference (teal = 15°C, salmon = 20°C) was detected. Black asterisk indicates significance (α = 0.05) between pre- and post-ash input. Shaded area denotes 95% confidence interval.

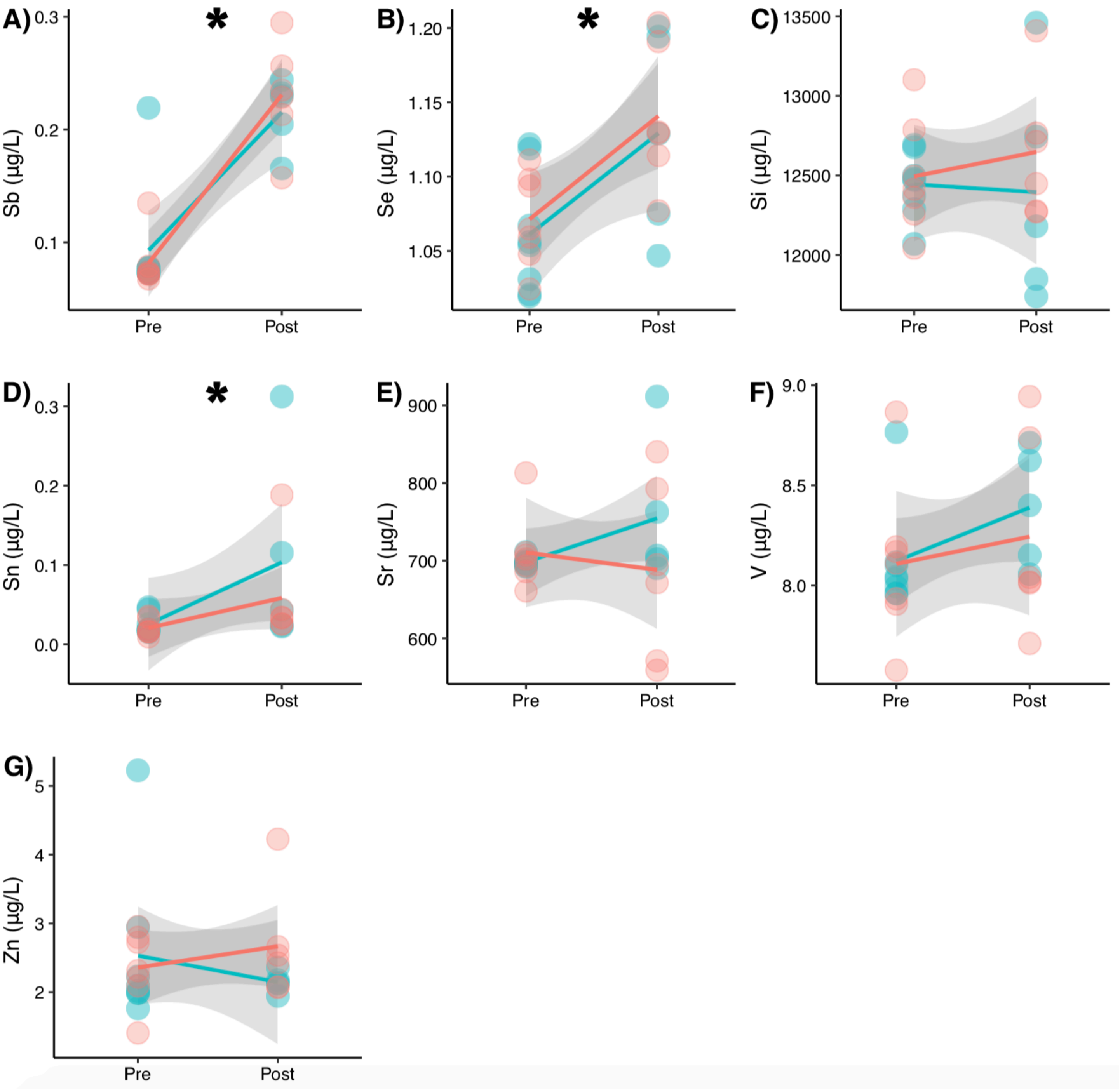


**Supplemental Figure 4:** Change in elemental concentration from ash input. Change in A) Sb, B) Se, C) Si, D) Sn, E) Sr, F) V and G) Zn concentration after ash input into experimental treatment water. No temperature difference (teal = 15°C, salmon = 20°C) was detected. Black asterisk indicates significance (α = 0.05) between pre- and post-ash input. Shaded area denotes 95% confidence interval.
